# The spatial proteome of the *Plasmodium falciparum* schizont illuminates the composition and evolutionary trajectories of its organelles

**DOI:** 10.1101/2025.11.27.690946

**Authors:** Scott A. Chisholm, Victor Flores, Alison Kemp, Lisa M. Breckels, Ludek Koreny, Nicolas Dos Santos Pacheco, Konstantin Barylyuk, Anna Kuroshchenkova, Kathryn S. Lilley, Julian C. Rayner, Ross F. Waller

**Affiliations:** Department of Biochemistry, University of Cambridge, Tennis Court Road, Cambridge CB2 1QW, UK; Cambridge Institute for Medical Research, University of Cambridge, Keith Peters Building, Hills Road, Cambridge, CB2 0XY, UK; Cambridge Centre for Proteomics, Department of Biochemistry, University of Cambridge, Tennis Court Road, Cambridge, CB2 1GA, UK

## Abstract

Malaria is caused by apicomplexan parasites of the genus *Plasmodium*, with all malaria symptoms and pathology caused by parasite stages that develop within, or transit between, host erythrocytes. The ability of *Plasmodium* cells to parasitise erythrocytes depends on distinctive intracellular compartments associated with invasion, as well as the development of unique cellular niches within the infected host cell. However, our understanding of the biology of the malaria parasite is limited by the fact that a large proportion of the parasite’s proteome has no known cellular location or function. To address this problem, we have generated comprehensive high-resolution maps of protein subcellular localisation for the invasive stage of the erythrocytic life cycle of *Plasmodium falciparum*, the major cause of malaria mortality. Using the spatial proteomics technique hyperplexed Localisation of Organelle Proteins by Isotopic Tagging (hyperLOPIT) we generated data for 3000 *P. falciparum* proteins expressed in late schizont stages. Our hyperLOPIT data resolved 24 distinct cellular niches, and using supervised machine-learning we can classify 1646 proteins into one of these compartments including exported sites within the host cell. Through comparative genomic analyses our data resolve the spatial patterns of cell evolution that have shaped the development of *Plasmodium* species and ongoing adaptive pressures and responses that challenge our efforts to manage these major disease-causing organisms.

## INTRODUCTION

Malaria remains a significant global health problem, particularly in Low- and Middle-Income Countries, with 249 million clinical cases and 621,000 deaths reported in 2023 (World Health Organization 2023). The causative agents of malaria are protistan parasites of the genus *Plasmodium*, with *P. falciparum* being the most lethal species responsible for almost all of the malaria-related morbidity and mortality. With both RTS,S/AS01 and R21/Matrix-M vaccines now being rolled out in endemic countries (Genton 2023), significant progress has been made in malaria prevention. Although these vaccines have shown moderate efficacy in reducing clinical malaria—most notably achieving around 70% protection in seasonal settings—they fall short of providing sterilising immunity or sustained protection across diverse parasite strains (Feehan, Plebanski, and Apostolopoulos 2025; Seidlein 2025). These limitations highlight the importance of continued research into parasite biology, including host–parasite interactions and immune evasion strategies, to guide the development of more durable interventions.

*Plasmodium* species are part of the phylum Apicomplexa, a diverse assemblage of obligate intracellular unicellular parasites that form specialized organelles and molecular structures at their anterior pole, collectively termed the apical complex, that play crucial roles in host cell invasion across the phylum (Waller and Carruthers 2024). Central to the apical complex are specialized secretory organelles, including rhoptries, micronemes, and dense granules, each contributing distinct molecular functions vital for the successful invasion and establishment of the parasite within its host cells (Carruthers and Sibley 1997; Counihan et al. 2013; de Koning-Ward et al. 2016). In addition to these secretory components, apicomplexan parasites exhibit further sophisticated cellular architectures, notably the inner membrane complex (IMC), a cytoskeletal feature comprising flattened membranous vesicles beneath the plasma membrane that supports cellular morphology and facilitates parasite motility and host interaction (Ferreira et al. 2020; Kono et al. 2012). Another hallmark of apicomplexan biology is the presence of the apicoplast, a plastid organelle originating through endosymbiosis of an alga. The apicoplast houses synthetic pathways for key metabolites (Ralph et al. 2004; Lim and McFadden 2010; Elahi and Prigge 2023; Bulloch et al. 2024), making it an attractive target for antiparasitic drug development and therapeutic intervention.

A major challenge with studying and understanding *Plasmodium* biology is interpreting both molecular divergence and novelty. As distantly related eukaryotes to most model systems many elements of common processes have diverged, even the relatively core eukaryotic machinery (Spielmann et al. 2020; Erath, Djuranovic, and Djuranovic 2019; Matthews, Duffy, and Merrick 2018; Yee et al. 2022; Dey and Baum 2021; Zeeshan et al. 2025; Schmidt et al. 2023). Furthermore, as parasites, they have responded to reductive evolutionary pressure for rapid growth and replication, resulting in genome reduction and much gene loss (Woo et al. 2015). Countering this, the challenges of combatting host defences systems and interacting with multiple host types and tissues mean they also must generate novel adaptations to be effective parasites. Indeed, large-scale *Plasmodium* genomic and population-genetic studies have catalogued remarkable variation of gene content across *Plasmodium* species and global parasite populations — including species-specific gene families, rapid evolution driven by host adaptation, and widespread diversity associated with drug resistance (Otto et al. 2018; MalariaGen et al. 2023; Wasmuth et al. 2009; Pearson et al. 2016; Miotto et al. 2024; Belda et al. 2025; Carlton et al. 2008). Therefore, for most *Plasmodium* genes, particularly those without orthologues in model organisms, we lack basic functional context, including where their protein products operate within the cell. To address this, we sought to provide a comprehensive subcellular framework for this novelty and divergence by determining the location of parasite proteins within the organelles, compartments, and niches where they act. We have focused on *P. falciparum* as the species most important to human health, and the invasion-ready schizont stage of the blood infection that represents the many adaptations relevant to the proliferation and expansion of these parasites within humans, as well as the pathology that they cause.

## RESULTS

### hyperLOPIT provides robust separation of subcellular compartments in late-stage *P. falciparum* schizonts

To determine the steady-state localisation of the proteome in *P. falciparum* mature schizonts we adapted the spatial proteomic method hyperplexed Localisation of Organelle Proteins by Isotopic Tagging (hyperLOPIT) (**Figure 1A-C**). The hyperLOPIT method has been successfully used for mapping the proteomes of other unicellular parasites: *Toxoplasma gondii, Cryptosporidium parvum* and two species of African trypanosomes (Barylyuk et al. 2020; Guerin et al. 2023; Moloney et al. 2023). In this method, cells are mechanically lysed using methods that largely preserve intracellular compartments intact, and these compartments are then dispersed on density gradients according to their different buoyant density properties (Mulvey et al. 2017). The abundance distribution profiles of all proteins across different fractions of these gradients are determined by multiplexed quantitative proteomics. Proteins with shared abundance profiles across the gradient provide evidence of protein co-location in the cell and, using markers of known cell compartments, novel or unknown proteins can be assigned to locations by machine learning (Crook et al. 2020; Crook et al. 2018; Christoforou et al. 2016; Hutchings et al. 2025).

**Figure 1.**
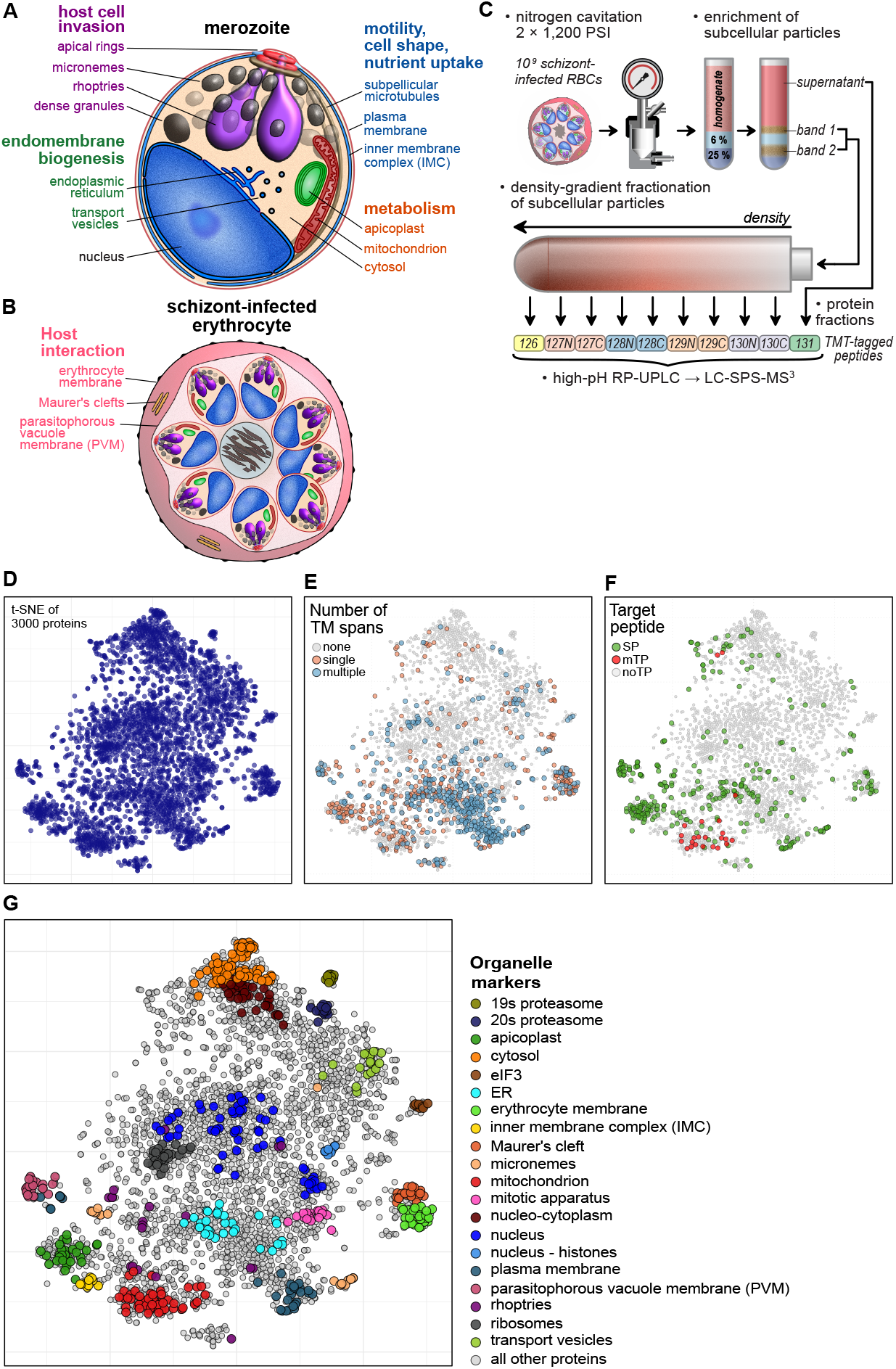
hyperLOPIT resolution of protein subcellular location gives cellular context to both conserved and novel proteins. **A**) and **B**) Schematics of *P. falciparum* merozoite and mature schizont within the erythrocyte showing main cell compartments **C**) Schematic of hyperLOPIT workflow. Cells are mechanically lysed, the homogenate is fractionated on density gradients, and gradient protein peptides tagged with unique TMT tags for relative quantitation by tandem mass spectrometry (LC-SPS-MS^3^) to generate abundance distribution profiles of each protein for correlation analysis as the basis of co-location. **D**) A 2D-projection of the abundance distribution profiles across 20 gradient fractions on 3,000 *P. falciparum* proteins (dots) shared across two hyperLOPIT datasets. t-distributed stochastic neighbour embedding (t-SNE) was used for dimensionality reduction, and clustering of proteins reflects similarity of their abundance distribution profiles. **E**) Predicted number of transmembrane (TM) spans for each protein shown on the t-SNE cell map. **F**) Predicted targeting peptides for protein entry into the endomembrane system (SP, Signal Peptide) or mitochondrion (mTP, mitochondrial Targeting Peptide). **G**) Position in the t-SNE projection of 542 curated marker proteins for known *P. falciparum* compartments or cellular niches.

We used nitrogen cavitation to disrupt schizont-stage *P. falciparum* given the success of this approach with other parasites. To enrich for organelles and less abundant compartment proteins, cell lysates were first centrifuged onto iodixanol density cushions. Protein from above the cushions were removed as a cytosol-enriched fraction, and the cytosol-depleted material at the cushion interfaces was further fractionated on iodixanol linear density gradients. Fractions taken across these gradients, and the cytosolic fraction, were labelled with unique tandem mass tags (TMTs) and the relative protein abundances across the gradients were determined by mass spectrometry. We performed two independent hyperLOPIT experiments on schizont-stage parasites (S1 & S2), adjusting cell preparation and ion concentration, to more comprehensively separate and resolve different subcellular niches. In experiment S1, we harvested schizonts tightly synchronized at the parasite ring-stage and allowed to progress to schizonts. In S2, we additionally used the PKG inhibitor Compound 2 (Baker et al. 2017) that prevents schizont egress and generates arrested mature schizonts.

From the two hyperLOPIT experiments, 3,581 proteins were fully quantified across 10 density gradient factions of one or the other experiment. Of these, 3,000 were common across both experiments, allowing the data from 20 fractions for 3,000 proteins to be concatenated for hyperLOPIT analysis. These multi-dimensional data were initially visualised using dimensionality reduction by t-distributed stochastic neighbour embedding (t-SNE) (Van der Maaten and Hinton 2008). These t-SNE projections indicated structure in these data with clusters of proteins indicating their common gradient distribution profiles (**Figure 1D**). To assess if this structure represented cell compartments and/or niches, we mapped the distribution of several general protein features that are indicative of protein function and location. Initial analysis revealed partitioning of proteins with predicted transmembrane domains indicative of segregation of cytosolic versus membrane associated cell features (**Figure 1E**). Predictors of signal peptides and mitochondrial targeting peptides (Almagro Armenteros et al. 2019) also showed multiple clusters enriched in one or other of these signals consistent with several clusters representing components of the endomembrane system and one putative mitochondrial cluster (**Figure 1F**).

To further test for cellular resolution in the *P. falciparum* hyperLOPIT map, and to assign cellular identities to protein clusters, we curated marker proteins of known organelles or niches drawing on the *Plasmodium* literature, predicted protein function, and predicted location (e.g., the ApiLoc database (Woodcroft et al. 2011)). We compiled these markers for conventional eukaryotic organelles, compartments and major complexes such as the nucleus, plasma membrane, mitochondrion, ER, cytoplasm, ribosomes and 26S proteasome components. Additionally, we compiled markers for known parasite-specific organelles such as the apicoplast, micronemes, rhoptries, and exported proteins that reside in the parasitophorous vacuole membrane (PVM), Maurer’s clefts, and the host cell membrane. Mapping these markers onto the t-SNE projections showed clear separation of subcellular components in our spatial proteome (**Figure 1G**), with the only exception being rhoptry marker proteins which were dispersed in this map (**Figure S1**, and addressed below). Further unassigned clusters were identified in the data either through using (i) unsupervised clustering of the untransformed abundance distribution profiles by hierarchical density-based spatial clustering of applications with noise (HDBSCAN) (Campello, Moulavi, and Sander 2013) (**Figure S2**), or (ii) by visual inspection of the t-SNE projections. Gene Ontology (GO) term enrichment analysis of four of these regions indicated proteins associated with the initiation factor eIF3, protein exchange between the nucleus and cytoplasm (‘nucleo-cytoplasm’), membrane and vesicular trafficking (‘transport vesicles’), and the mitotic apparatus (**Figure 1G** and **Figure S3**) and these proteins were subsequently added as markers for these further clusters. Overall, we compiled a list of 542 marker proteins which indicated clear segmentation of different subcellular compartments with obvious identities, including resolution of compartments within the host erythrocyte such as the PVM and Maurer’s clefts.

The lack of a resolved rhoptry organelle proteome in our initial spatial proteome was speculated to be due to the dual location of many rhoptry proteins in the intra-erythrocyte schizont stage. In this stage, rhoptry proteins would be simultaneously present both within new rhoptries of merozoites ready for the next invasion cycle, and as previously secreted proteins from the current invasion cycle where they played key roles in establishing the infection. Examples of these secreted rhoptry proteins destinations include components of the *Plasmodium* surface anion channel (PSAC) in the erythrocyte membrane, and RAP proteins at the PVM (Counihan et al. 2017; Nguitragool et al. 2011; Sherling et al. 2017). To untangle these signals and distinguish resident, rhoptry proteins from exported rhoptry proteins, we generated a third hyperLOPIT dataset (S3) from purified merozoites where exported proteins should be depleted (Table S1). We treated schizont-stage parasites with the cysteine protease inhibitor E-64 that allows parasite disruption of the PVM but blocks egress from the erythrocyte. This enables mechanical release and isolation of merozoites prior to lysis by nitrogen cavitation (Boyle et al. 2010). From these merozoite-enriched samples we generated hyperLOPIT protein abundance profiles that represented 2753 proteins, with 2496 proteins common with the two previous schizont hyperLOPIT datasets (S1 & S2) (**Figure 2A**). The combined data from these three experiments (S1-S2-S3), when viewed as t-SNE projections, showed the rhoptry marker proteins as a single distinct cluster, indicating the approach of depleting host cell membranes in the merozoite sample mitigated some of the issues observed with the rhoptry proteins of the schizont samples alone (**Figure 2B**).

**Figure 2.**
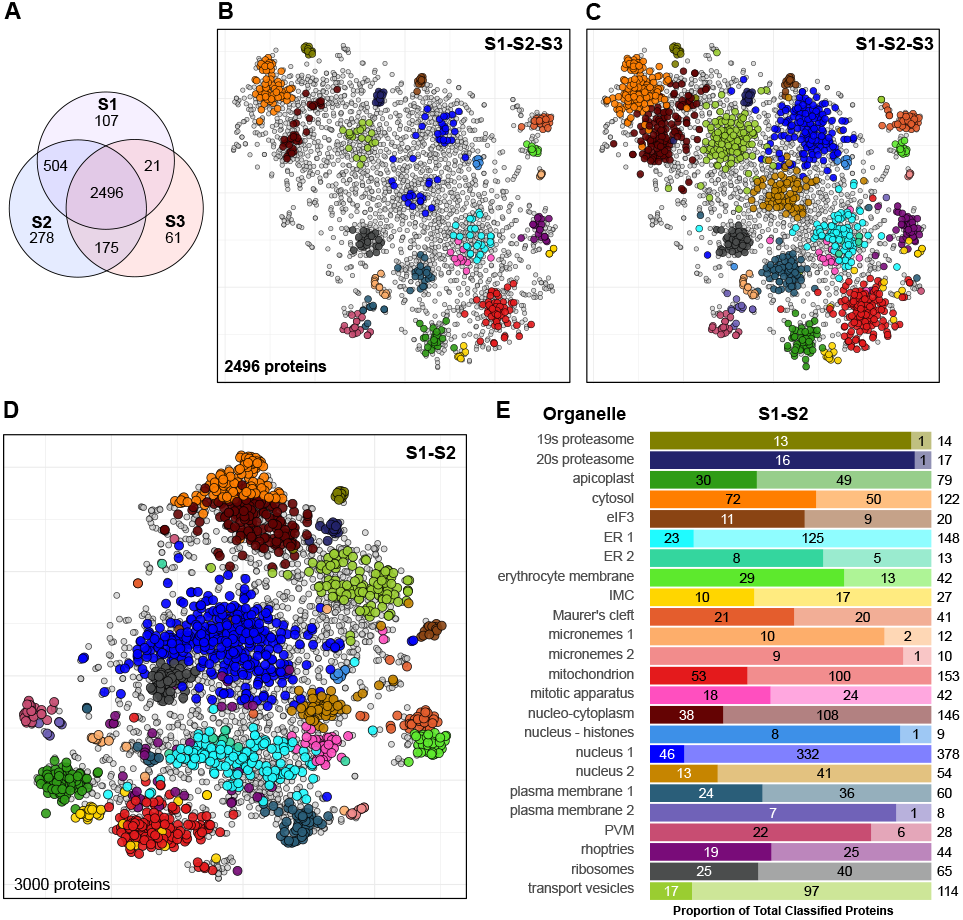
Protein assignment to *P. falciparum* subcellular niches from three hyperLOPIT datasets. **A**) Venn diagram showing the overlap of proteins quantified across three hyperLOPIT experiments: S1-S2-S3. **B**) A t-SNE projection of the abundance distribution profiles of 2496 proteins (dots) common to all three datasets (S1-S2-S3) showing compartment markers (coloured dots) used as training data for protein localisation prediction. **C**) A t-SNE projection showing the assignment of proteins by colour to one of 24 compartments following localisation prediction using a Support Vector Machine (SVM) across the three datasets (S1-S2-S3). Grey points represent unassigned proteins. **D**) SVM assignment of proteins to one of 23 compartments (minus rhoptries) across two datasets (S1-S2). **E**) Barplot showing the number of proteins assigned to each compartment from the S1-S2 analysis (left, solid colours) and predicted proteins (right, opaque colours). Rhoptry assignment numbers are taken from the S1-S2-S3 analysis (C). Niche protein totals are on the far right. Colours indicating compartment in E) are also used in B-D).

### Classification of unknown proteins to subcellular niches

Using the abundance distribution profiles of the markers as labelled training data for supervised machine learning, we classified proteins of unknown localisations to parasite subcellular niches. A Support Vector Machine (SVM) was used to define the boundaries in the untransformed data around each niche using the R package pRoloc (v1.48.0)(Gatto et al. 2014). Some marker sets were observed to project as two distinct clusters on the t-SNE projections (micronemes, plasma membrane, ER, nucleus) (**Figure 1G, 2B**) and visual inspection of the abundance distribution profiles support this distinction (**Figure S4**). We, therefore, separate these four niches each into two marker populations for the assignment of unknown proteins. Protein localisation prediction revealed 1375 new protein assignments across 24 compartments, of the 2496 proteins common in the S1-S2-S3 data (**Figure 2C, S5**). This included 25 unknown proteins assigned to the rhoptry proteome totalling 44 rhoptry proteins. Given that the combined S1-S2 schizont data contained over 500 additional common proteins, we also applied SVM classification to these data, but without the rhoptry markers given their lack of resolution in these samples. This analysis yielded a further 229 proteins assigned to one of the 23 niches, and incorporating the rhoptry classified proteins from the three-way analysis, gave 1646 proteins classified to one of 24 subcellular niches (**Figure 2D, E, Table S2**).

To test the robustness of the hyperLOPIT clusters, 15 uncharacterised proteins associated with distinct subcellular clusters were selected for reporter-tagging by endogenous C-terminal HA epitope-fusion using a modified selection-linked genome integration system (Birnbaum et al. 2017; Jonsdottir et al. 2021)(**Table S3**). Of the 15 proteins attempted, drug-resistant parasites could be recovered for all after G418 selection. Of these, 11 showed correct genomic integration of the tagging construct by diagnostic PCR, and Western blotting detected an expressed band of the expected molecular weight (**Figure S6**). HA signal was also detectable by immunofluorescence microscopy assays (IFA) for 9 of these candidates, all of which localised in accordance with their predicted hyperLOPIT clusters (**Figure 3**). Together, these results provide experimental validation that the hyperLOPIT location assignments faithfully capture organelle-specific protein distribution in *P. falciparum*.

**Figure 3.**
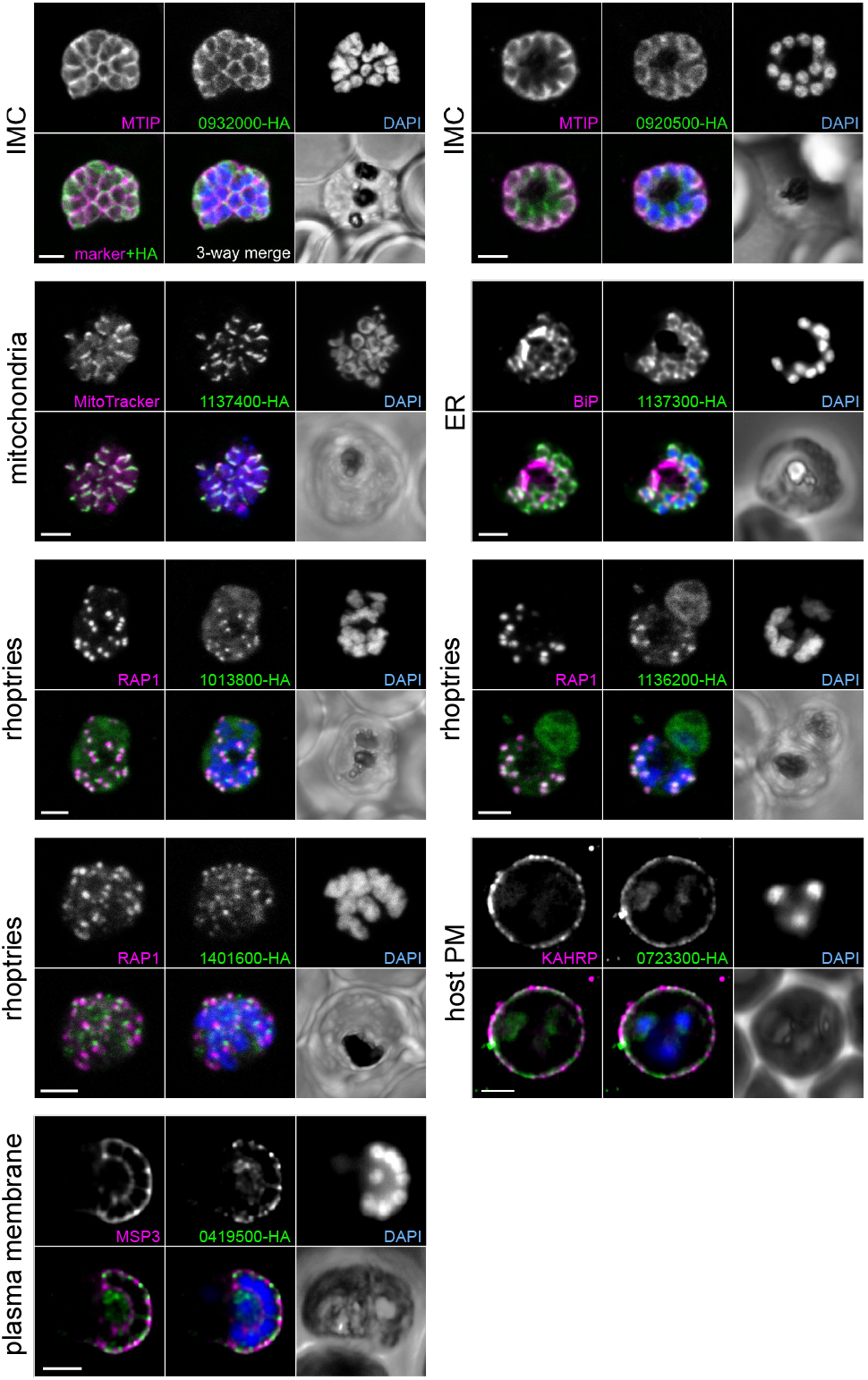
Validation of hyperLOPIT-predicted subcellular locations. Selected uncharacterised proteins were endogenously 3xHA-epitope tagged and detected by immunofluorescence assays (green) and co-stained for known organelle markers (magenta). DNA stained with DAPI (blue) and merges of HA+marker and HA+marker+DNA are shown in the lower panels in addition to phase imaging of the infected erythrocyte. Scale bar = 2 µm.

### Spatial proteomics reveals cellular distribution of protein novelty, reduction, selection and adaptation in *Plasmodium* parasites

The clustering of proteins by hyperLOPIT into cohorts sharing distinct biochemical niches provides an unprecedented model of the location of the proteome of the late schizont and the blood-invasive merozoite of *P. falciparum* parasites. With this information, we can assess the biochemical and evolutionary characteristics of different compartments to better understand the molecular evolution, adaptation and biology of this parasite.

### How has gene gain and loss driven *Plasmodium* evolution?

Apicomplexans are conspicuous for their abundance of novel gene families, and the fixation of new genes and/or the expansion of gene families is one mechanism that drives adaptation in parasites (and all life). The location of the products of new *Plasmodium* genes, and the evolutionary timing of their origin or expansion, reflects which facets of parasite biology that have been most active in driving *Plasmodium* evolution. To determine the evolutionary timing of new gene origins in *P. falciparum*’s past, we conducted gene orthology analysis of 21 available genomes across Apicomplexa, including multiple *Plasmodium* species, haemosporidians, hematozoans and coccidians. This analysis identified when novel gene orthogroups originated in each phylogenetic clade and species (**Figure 4A**). At each phylogenetic level, we then examined the distribution of new proteins across subcellular compartments to test if there were any biases reflecting pronounced compartment evolution at a given evolutionary time compared to the rest of the cell—the subcellular choreography of parasite evolution (**Figure 4B**). This analysis showed that recently developed orthogroups that are unique to *P. falciparum*, and that are detectable in the late schizont stage, are strongly enriched in the parasite plasma membrane, as well as those exported to the Maurer’s clefts and erythrocyte plasma membrane. This evolutionary trend represents a consistent and deeper hallmark of *Plasmodium* evolution, with equivalent innovation in these compartments, as well as in the parasitophorous vacuole membrane, seen through the radiation of the Laverania subgenus that infect African apes and from which *P. falciparum* evolved (Rayner et al. 2011), as well as extending back to the diversification of many mammal-infective *Plasmodium* spp. Proteomic novelty gained in the common ancestor of the *Plasmodium* genus, however, is much more widely spread throughout cell compartments including the primary structures of invasion (micronemes, rhoptries and IMC) as well as general metabolic and cell regulatory compartments. These data indicate that broad cell compartment remodelling through the acquisition of new proteins preceded the origin of the *Plasmodium* genus, but that subsequent parasite adaptations to different hosts has seen this novelty concentrated at the interfaces of interaction between parasite and host after erythrocyte invasion has occurred.

**Figure 4.**
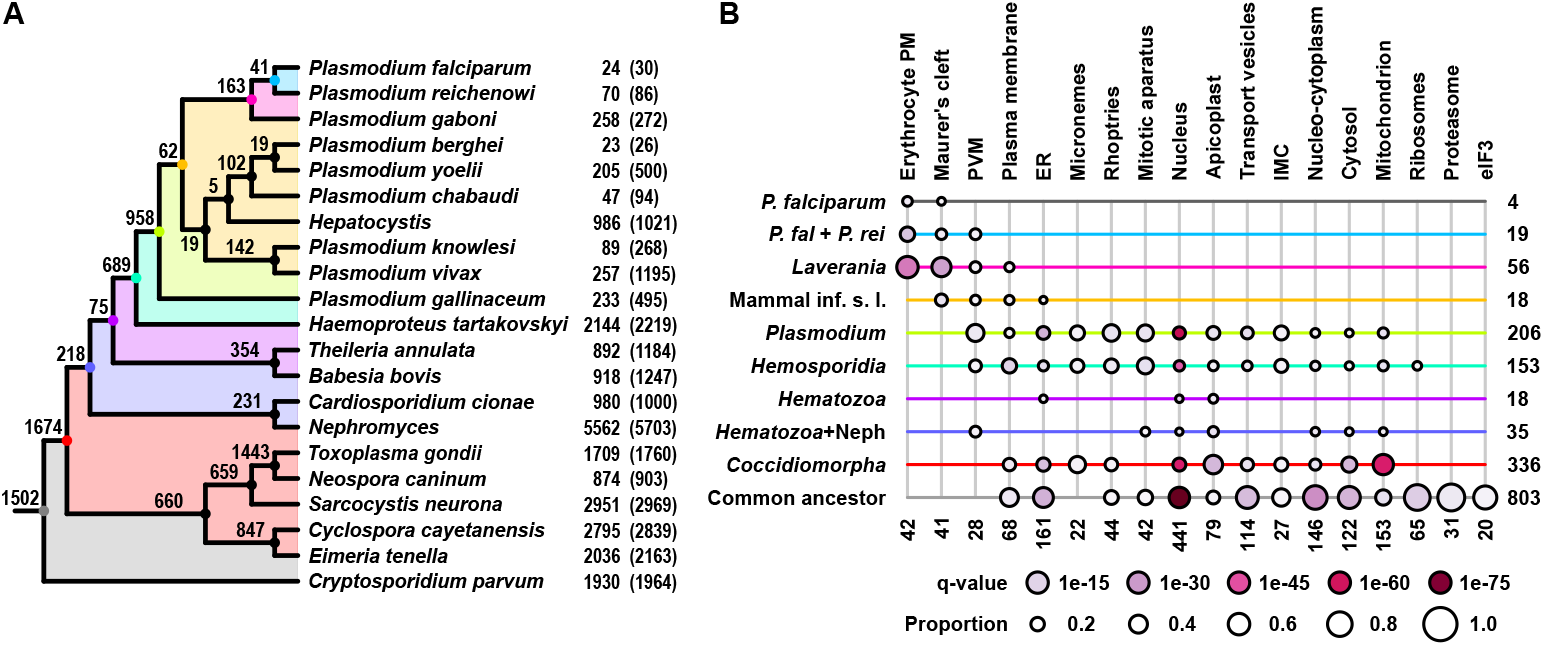
Subcellular distribution of new protein orthogroups over evolutionary time. **A**) Numbers of novel protein orthogroups to Apicomplexa shown according to when in apicomplexan phylogeny they arose. The number of orthogroups unique to each species is shown to the right of species names, and numbers in brackets denote unique proteins. **B**) Distribution of significant enrichments for new protein orthogroups by subcellular location for each phylogenetic position leading to *P. falciparum*. Cumulative orthogroup protein numbers are shown for each phylogenetic level (right) or subcellular compartment (bottom). Significance is shown as q-values, and proportion indicates the fraction of the total subcellular compartment proteome found in *P. falciparum*. “Common ancestor” represents the genes common to the apicomplexan common ancestor.

Parasitism typically entails strong selective pressure for genome reduction, and this is particularly evident in *P. falciparum* whose genome is small even compared to other apicomplexans (Gardner et al. 2002; Swapna and Parkinson 2017). In this context of reductive selective pressure on the diversity of the protein components of the cell, we have asked which cell components show evidence of greatest reduction to a minimal essential proteome, versus those where multiplicity of functional proteins is required. As a measure for essential protein function during *P. falciparum* asexual *in vitro* propagating, we have used the data from a saturation mutagenesis screen (Zhang et al. 2018) where relative growth associated with gene disruption was used to attribute a ‘mutagenesis fitness’ score (MFS) for all *P. falciparum* genes (lower scores indicating genes less amenable to disruption). Our subcellular proteome enables the distribution of MFS values for each functional cell compartment to be assessed (**Figure 5**). These data show that the apicoplast, transport vesicles, ER and ribosomes are the most significantly enriched for proteins essential for the asexual growth cycle. By contrast, the proteomes of cell compartments that directly interact with the host (erythrocyte plasma membrane, Maurer’s clefts, PVM, parasite plasma membrane) including during invasion (micronemes) are all biased towards proteins with less or no phenotypes when disrupted in the *in vitro* propagation context. This is also the case for the nuclear proteome as a whole (although excluding the nucleus-cytoplasmic exchange functions). Collectively these data suggest that greater diversity of proteins has been prioritised in host interactions and nuclear regulation in this context of gene loss and genome reduction.

**Figure 5.**
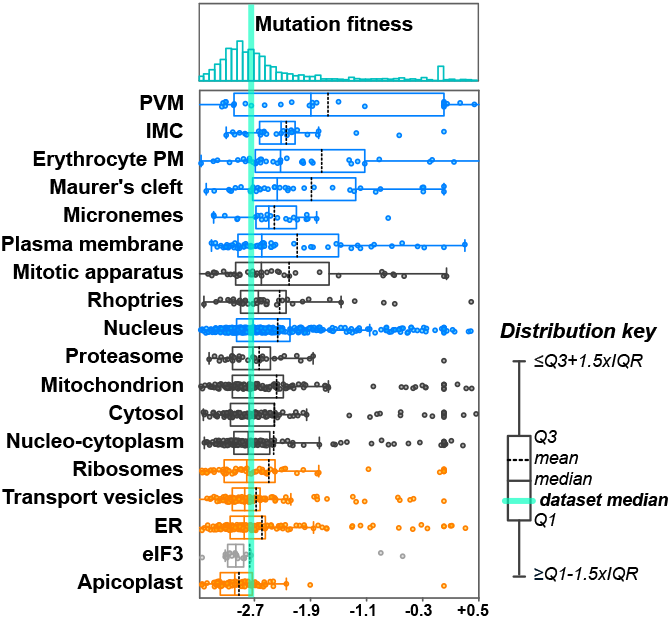
Distribution of Mutagenesis Fitness Scores (MFS) across cell compartments for schizont-expressed proteins. Cell-wide distribution of MFS is shown in the histogram (above) and distributions of MFS for each compartment are shown for individual proteins (dots) and overall as box plots (legend at right). Compartments are ranked according to median values, and those significantly enriched for proteins with low MFSs (more essential proteins) are shown in orange, and for higher MFSs shown in blue. Distributions in black showed no significant differences from the average cell-wide median MFS value, and distributions in grey lacked sufficient proteins for significance testing. Q1, first quartile; Q3, third quartile; IQR, inter-quartile range.

### Where in the cell do mutations and selection drive intra and inter-species adaptations?

Mutations of existing genes is also a critical driver of organism evolution, and the distribution of gene mutation density across different compartment proteins can indicate where this process is most affecting change in these parasites. *P. falciparum* genomic diversity has been captured within the MalariaGen project representing over twenty thousand clinical *P. falciparum* isolates from across the world (MalariaGen et al. 2023). Mapping gene mutation density onto our compartment proteomes shows a clear enrichment for mutations in genes associated with host-interactions (**Figure 6A**). This bias is seen for all the surface or extra-parasite sites in the infected erythrocyte, as well as the micronemes and rhoptries. The nucleus and mitotic apparatus are also prone to high mutation densities; however, most metabolic compartments (cytosol, apicoplast, mitochondria and ribosomes) are biased for lower accumulation of mutations. Positive selection for change is also evident from elevated pN/pS ratios—the ratio of non-synonymous to synonymous mutations rates within a population where higher values indicate adaptive evolution. Enrichment for high pN/pS values is again seen at all sites that mediate host interaction and showing parasite responses to this strong positive selection for change (**Figure 6B**). Compartments under strongest purifying selection biased for cytosolic functions and the mitochondrion.

**Figure 6.**
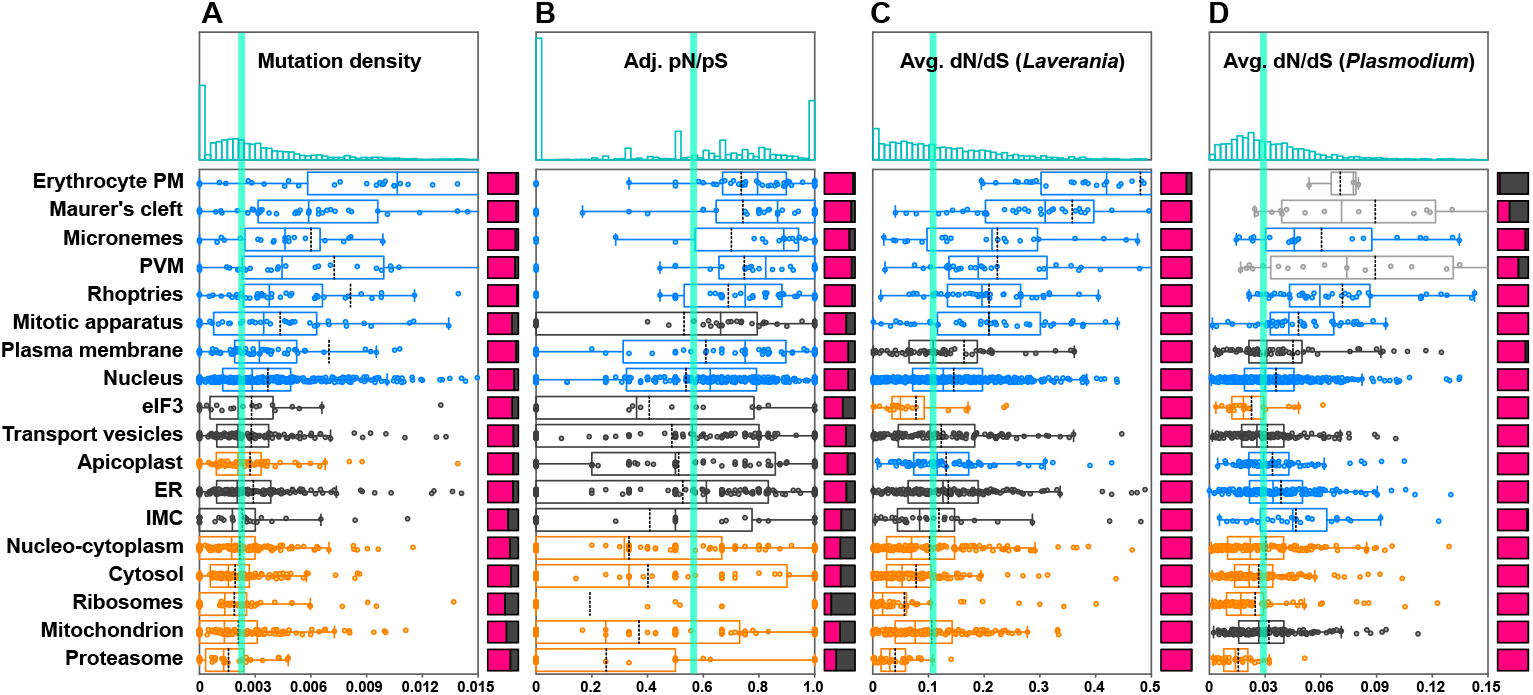
Distribution of mutational and selective pressures on *Plasmodium* compartments. Compartment-specific distributions of **A**) gene mutation density as single nucleotide polymorphisms per nucleotide, **B**) within *P. falciparum* population selective pressures expressed as protein average adjusted pN/pS, **C**) within *Laverania* selective pressure expressed as protein average dN/dS, and D) within *Plasmodium* genus selective pressure expressed as protein average dN/dS. Distributions for the whole cell are shown as histograms (above) and for each compartment as box plots with statistical skews to higher (blue) or lower (orange) values as for Figure 5. Compartments are ranked by median values for A). Red/grey bars to the right of each profile show either the proportion of proteins with zero values (grey) (A and B), or the proportion of *P. falciparum* proteins with orthologues (red) in all © or >4 (D) of the species being compared.

Speciation in *Plasmodium* has often accompanied expansion into new vertebrate hosts, and between species average rates of non-synonymous to synonymous mutations (dN/dS) show where positive selection has been instrumental in adaptation to these changed environments. Using the *P. falciparum* spatial proteome as a reference for the whole genus, we have asked where in the cell positive selection has been strongest during speciation versus within a species population diversity (*P. falciparum*). Within the *Laverania* group the sites mediating host interaction, again, are under strong selection for change between Species (**Figure 6C**). However, positive selection is also now seen on the apicoplast as well as the mitotic apparatus. These trends are also seen across all 10 species examined (including *Hepatocystis*) although, in addition to the apicoplast and mitotic apparatus, positive selection is also seen on the IMC and ER driving speciation processes in *Plasmodium* (**Figure 6D**).

### Where do biases for protein disorder and size occur in the cell?

Generic properties of a protein such as intrinsic disorder and size often influence their functionality and evolutionary flexibility, as changes in these features can facilitate the emergence of novel interactions or functions. We therefore tested whether there were biases in the distribution of these protein features in schizont-stage *P. falciparum*. We used multiple predictors of protein disorder: IUPred3 (Erdos, Pajkos, and Dosztanyi 2021), MobiDB-lite (Necci et al. 2017) and two outputs from Alphafold, DSSP and confidence scores (Varadi et al. 2024). We assigned disordered regions to any protein where there was consensus of two or more methods for any intrinsic disordered domain (IDD) ≥30 residues, and expressed IDD scores the disordered length fraction of the total protein length. Most cell proteins lacked IDDs, and cytosolic and metabolic compartments were generally most depleted in IDDs (**Figure 7A**). Compartments that showed positive skews towards proteins with high disordered fractions were the nucleus, including the mitotic apparatus, and the PVM and Maurer’s cleft export compartments. Protein size also showed non-uniform distribution across cell compartments (**Figure 7B**). Again, the nucleus and mitotic apparatus are strongly biased for large proteins suggesting that size and disorder correlate here. The rhoptries also show a bias for large proteins, whereas the PVM and Maurer’s clefts are skewed to smaller than average proteins suggesting that small, disordered proteins are important constituents of both compartments.

**Figure 7.**
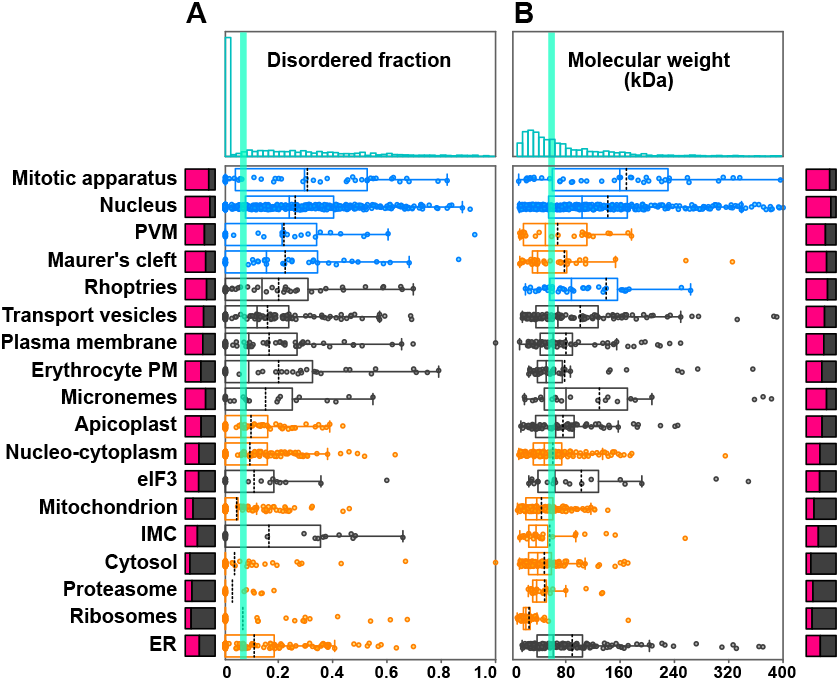
Compartment distributions of protein disorder and molecular weight. **A**) Distributions of predicted disordered protein regions as a fraction of the total protein length for each compartment. Bars to the left show in red the proportion of proteins with any contiguous predicted disordered region ≥30 amino acids. Compartments ranked on median values. **B**) Distribution of protein molecular weights in kDa by compartment. Distributions for the whole cell are shown as histograms (above) and for each compartment as box plots with statistical skews to higher (blue) or lower (orange) values as for Figure 5.

## DISCUSSION

The hyperLOPIT spatial proteomic method has enabled a high-resolution map of protein locations in the late-stage *P. falciparum* schizont for the first time. Moreover, by performing this technique on parasites contained within the infected erythrocyte where the parasite has established an elaborately modified niche, we have also resolved compartments beyond the parasite containing exported *P. falciparum* proteins. These sites include the parasitophorous vacuole, Maurer’s clefts and erythrocyte plasma membrane providing further insight into the parasite’s interaction with the host. A challenge for the hyperLOPIT approach on parasites within their hosts, however, is the potential for secreted proteins to occupy multiple locations: their pre-secretion compartments and post-secretion destinations which will disrupt their resolution as a single organelle cluster. This was the case for rhoptry proteins from the first two schizont samples which failed to resolve as one cluster, but did show some evidence of clustering of proteins of common final destination (e.g. the RAP proteins that reside in the parasitophorous vacuole, and the CLAG/RhopH complex whose final destination is the erythrocyte membrane). By generating separate merozoite-enriched samples these confounding multiple signals were reduced sufficiently that the rhoptry organelle could be resolved and a cohort, and 44 rhoptry proteins were identified. By contrast, the PVM was resolved as one cluster but is known to be populated by different parasite secretory vesicles: both dense granules and post-invasion constitutive secretion routes (Bullen et al. 2012; Richard et al. 2009; Riglar et al. 2011). Therefore, these maps must be interpreted as steady-state distributions of proteins in the context of their biological state.

This *P. falciparum* map shares many features with similar maps for other apicomplexans such as *Toxoplasma gondii*. However, there are some compartments that resolve better in one parasite over the other. For instance, the conoid and other structures of the apical complex resolved well in *T. gondii*, but we found that the *Plasmodium* orthologues of this structure were not resolved in our corresponding maps. This likely reflects the differences in stability or robustness of different structures between different species, which could also be the case between cell stages of a single species. The *T. gondii* tachyzoite conoid is known to be a robust structure tethered to an elaborate IMC and subpellicular microtubule cage. In contrast, the *P. falciparum* merozoite lacks a tubulin conoid structure and has a greatly reduced structural component of the IMC (only two subpellicular microtubules (Bannister et al. 2003; Ferreira et al. 2023)). These structures likely dissociated during the sample lysis conditions used. On the other hand, a cluster enriched in elements of the mitotic machinery is evident in the *P. falciparum* maps but was not in the *T. gondii* extracellular tachyzoites. This could reflect differences in cell stage, with some elements of this post-division apparatus still presented in our schizont samples, but perhaps little cell division represented in the *T. gondii* samples. Similarly, in *P. falciparum* we see microneme proteins resolving as two clusters, consistent with a recent study using ultrastructure expansion microscopy, where AMA1 and EBA175 were seen in distinct microneme populations (Liffner et al. 2023). It could be that differences in physical properties of microneme types would enable hyperLOPIT resolution. We note, however, that these two populations of microneme proteins were only obviously diverged in their abundance distribution profiles in the S1 sample (Fig. S3) where the non-arrested schizonts were likely not as tightly synchronised as the Compound 2-treated S2 sample, nor the release E64-treated merozoites. Thus, the *microneme1-2* separation might also indicate a developmental difference in either the timing of synthesis or maturation of these two populations. In any case, *microneme2* is enriched in proteins associated with early events of merozoite attachment to the erythrocyte, and *microneme1* for invasion events, suggesting at least some level of separation of delivery of these functional elements of invasion.

Collectively, by combining the signals from three samples representing the maturation of *P. falciparum* schizonts, we give context to 1646 proteins that, importantly, includes many of the novel and uncharacterised proteins of these parasites. Some 24 subcellular niches in total were resolved which, in turn, provides a framework for more fully investigating the functions of the compartmentalisation of these cells. Further definition of compartment proteomes can be limited either by a sufficient number of positive markers or by our knowledge of protein associations and the functional compartments or niches that they represent. The observed resolution of some small functional clusters (e.g. histones) indicates that there is further signal for protein association within the spatial maps of these proteins that awaits discovery and investigation.

The generation of an objective sample of organelle and compartment proteomes of the schizont, even in the absence of knowledge of the function of all of these proteins, enables them to be used as a measure of compartment role, selective pressure and evolutionary trajectory in this important stage of pathogen propagation. This provides both new insights and revisions of previous interpretations. The apicoplast, for example, while known to contain pathways for multiple anabolic processes, has been understood to only be required for the synthesis of isopentenyl pyrophosphate (IPP) for isoprenoids in blood-stage parasites(Yeh and DeRisi 2011; Gisselberg et al. 2013). However, knockout phenotypes from genome-wide mutagenesis screens shows that virtually all detectable elements of the schizont-stage apicoplast proteome confer strong phenotypes. We note that while some of the proteins known to be part of the pathways for fatty acid synthesis and tetrapyrrole synthesis for haem were not detected in our schizont proteomic data, even these have strong blood-stage phenotypes when disrupted, suggesting likely low expression levels but nonetheless provision of essential function for parasite propagation. The growth of synthetic apicoplast-lacking cells in IPP-supplemented conditions might be explained by persistent function of expressed apicoplast proteins even without an intact organelle, or phenotype levels below the detection limits of these supplemented conditions (Yeh and DeRisi 2011; Gisselberg et al. 2013). Similarly, predictions of blood-stage dispensable apicoplast genes made from the genome-wide mutagenesis screen (Zhang et al. 2018) could indicate genes for apicoplast proteins that are not expressed in these parts of the lifecycle, or false-positive predictions of apicoplast location. The importance of apicoplast function in *P. falciparum* is further indicated by its skewed for lower accumulation of mutations in its protein-encoding genes compared to the rest of the cell. Surprisingly, however, at a speciation level there is evidence of positive selection for change in apicoplast proteins that is not seen within intra-specific populations of *P. falciparum* itself. While it is difficult to predict the impact of these change to apicoplast proteins on organelle function, these data suggest that elevated change within this organelle has accompanied changes of host species. As a metabolic organelle, this could reflect differing balances between scavenging and self-sufficiency for some key metabolites required in the different host species environments. Apicoplast evolution is also evident prior to the origin of *Plasmodium* species with novel extant *P. falciparum* proteins consistently recruited to this organelle throughout the apicomplexan radiation. Thus, the complex yet essential role of the apicoplast in *Plasmodium* biology is revealed by its proteome which continues to highlight it as a promising target for further drug-based treatments.

The *P. falciparum* schizont spatial proteome also illuminates the evolutionary importance of the proteome dedicated to the parasite’s interaction with the host. Novel proteins that have arisen within *Plasmodium* radiation are overwhelmingly deployed to the erythrocyte plasma membrane, Maurer’s clefts, PVM and parasite plasma membrane. This is also where the greatest apparent multiplicity of function is required (as judged by *in vitro* culture condition protein redundancies). Moreover, along with microneme and rhoptry proteins, this is where the signal for strongest selection for change is seen both within *P. falciparum* populations and during the radiation of the genus. It is expected that proteins that function while exposed directly to the host’s extracellular immune machineries are under strong adaptive evolution, however, these elevated dN/dS signals for Maurer’s cleft and PVM proteins are more surprising given their internal location within the erythrocyte. The Maurer’s cleft and PVM proteins are also enriched for disordered domains, a feature that has been suggested could contribute to immune evasion (Guy et al. 2015). Together, these features might indicate that the proteins of these compartments could be more relevant to the host immunity than previously expected. Disordered protein regions are also often linked to protein–protein interactions. The *Plasmodium* export pathways are not well understood (Levray et al. 2023; Beck and Ho 2021; Das 2025) but it is possible that the high degree of disorder in the proteins of these export routes allows for flexible, transient interactions during trafficking and/or host engagement. The relatively small size of the disordered proteins of the PVM and Maurer’s clefts may help to limit the risk of aggregation as the transit parasite and host compartments.

Other targeted studies seeking definition of *Plasmodium* compartment proteomes have been undertaken, including using sequential cell compartment lysis to resolve secreted proteomes (Siau et al. 2023), proximity-dependent biotinylation for apicoplast proteins (Boucher et al. 2018), and integrated computational approaches for mitochondrial proteins (van Esveld et al. 2021). Comparison of these data with our hyperLOPIT maps show broad overlap of prediction but, unsurprisingly, some disagreement also. These differences reinforce the importance of orthogonal approaches to understanding the organisations of these cells, and will necessarily include further approaches, on different cell stages, and in different conditions. Nevertheless, our data provide a substantial advancement to how we can study and interpret these cells with the ultimate goal of advancing our approaches to managing their impact on the global health burden that they are responsible for.

## ACKNOWLEDGEMENTS

Anti-RAP1 and anti-KAHRP antibodies were obtained from the European Malaria Reagent Repository (EMRR: www.malariaresearch.eu). Anti-PfGRP78 (BiP) antibodies were obtained through BEI Resources, NIAID, NIH. This work was supported by the Wellcome Trust, United Kingdom, Investigator Award 214298/Z/18/Z to R.F.W and Investigator Award 220266/Z/20/Z to J.C.R., and the Gordon and Betty Moore Foundation to R.F.W. (https://doi.org/10.37807/GBMF9194). L.M.B was supported by BBSRC strategic Longer and Larger (sLoLa) grant BB/T002182/1. N.D.S.P was supported by a “Postdoc.Mobility” fellowship from the Swiss National Science Foundation (P500PB_217798). We would like to thank Robin Antrobus for mass spectrometry processing, and we thank Olivia Carmo for useful discussions.

## AUTHOR CONTRIBUTIONS

S.A.C., K.S.L., J.C.R. and R.F.W. conceived this study. S.A.C performed and analysed the hyperLOPIT experiments, with V.F., L.M.B. and K.B. contributing to the design and implementation of the analyses. A. Kemp, L.K., N.D.S.P. and A. Kuroshchenkova performed gene-tagging location validations. S.A.C. and R.F.W. wrote the manuscript, which was read and approved by all authors.

## DECLARATION OF INTERESTS

The authors declare no competing interests.

## DATA AVAILABLITY

The mass spectrometry proteomics data have been deposited to the ProteomeXchange Consortium via the PRIDE (Perez-Riverol et al. 2025) partner repository with the dataset identifier PXD070842 and 10.6019/PXD070842. The data are integrated into PlasmoDB. An interactive interface to the annotated spatial proteome data is available via a web-based R Shiny application at https://proteome.shinyapps.io/plasmolopitsz/ (S1-S2) and https://proteome.shinyapps.io/plasmolopitszmz (S1-S2-S3).

## METHODS

### Contact for Reagent and Resource Sharing

Further information and requests for reagents may be directed to and will be fulfilled by the corresponding author Ross F. Waller.

### *Plasmodium falciparum* culture

*Plasmodium falciparum* (3D7 strain) parasites were maintained in human erythrocytes sourced from NHS Blood and Transplant (Cambridge, UK), with approval from the NHS Cambridge South Research Ethics Committee (20/EE/0100) and the University of Cambridge Human Biology Research Ethics Committee (HBREC.2019.40). Cultures were grown at 4% haematocrit in RPMI 1640 medium (Gibco, UK) supplemented with 5 g l−^1^ Albumax II, 2 g l−^1^ anhydrous dextrose EP, 5.96 g l−^1^ HEPES, 0.3 g l−^1^ sodium bicarbonate EP and 0.05 g l−^1^ hypoxanthine, dissolved in 2 M NaOH. Parasites were incubated at 37 °C in a gassed incubator under a low-oxygen atmosphere (1% O_2_, 3% CO_2_ and 96% N_2_; BOC, Guildford, UK).

### Synchronisation of parasites

Parasite cultures were synchronized using sorbitol treatment as previously described (Lambros and Vanderberg 1979). Briefly, ring-stage parasites were pelleted by centrifugation, resuspended in ten volumes of 5% D-sorbitol (Sigma-Aldrich) and incubated at 37 °C for 10 min to selectively lyse trophozoite- and schizont-infected erythrocytes. Cells were collected by centrifugation, washed once with RPMI 1640 and returned to culture at 4% haematocrit.

To further improve synchronization, Giemsa-stained smears were examined to confirm a predominance of schizont-stage parasites. The medium was removed, and infected erythrocytes were gently resuspended in 5 ml culture medium and layered over 10 ml 63% Percoll (Merck, P4937) prepared with 10% 10× PBS and 27% RPMI 1640. Gradients were centrifuged at 1,100 × g for 11 min with no brake and acceleration set to 3. The late-stage parasite band at the interface was collected, washed in medium and transferred to fresh uninfected erythrocytes at 4% haematocrit. Cultures were incubated at 37 °C in a gassed incubator for 3 h before a second round of sorbitol treatment as above.

In selected experiments, egress inhibitors were used to increase synchronicity at harvest. In experiment S2, parasites were treated with 1.5 µM Compound 2 (a PKG inhibitor) for 3 h after Percoll purification. In experiment S3, parasites were treated with 10 µM E-64 (a cysteine protease inhibitor) for 4 h (Boyle et al. 2010) followed by mechanical filtration through a 1.2 µm membrane filter to isolate merozoites for hyperLOPIT processing.

### Spatial Proteomics: hyperLOPIT

For each hyperLOPIT experiment, approximately 10^9^ Percoll-enriched schizont-stage parasites were harvested using Percoll as described above, washed once with ice-cold PBS (pH 7.4) and resuspended in 5 ml of chilled homogenisation buffer (0.25 M sucrose, 10 mM HEPES-KOH, pH 7.4, 2 mM EGTA) supplemented with EDTA-free protease inhibitors (Roche) as described in (Mulvey et al. 2017). In experiment S3, the E-64-arrested schizonts were resuspended in homogenisation buffer prior to passing through the 1.2 µm membrane filter.

Parasites were mechanically lysed by nitrogen cavitation (Hunter and Commerford 1961; Simpson 2010) using a 45 ml cell disruption vessel (Parr Instruments, model 4639) pressurised to 1,200 psi. The pressurised suspension was equilibrated on ice for 20 min with gentle agitation, then released through the outlet valve at a controlled rate of ∼2 drops per second. Intact or incompletely lysed cells were pelleted by differential centrifugation and subjected to a second cavitation cycle. Supernatants from both cycles were pooled to generate the crude cell homogenate. To reduce viscosity, the homogenate was treated with 500 U Benzonase nuclease (Millipore, 71205) for 20 minutes at room temperature, followed by a further 10 minutes on ice.

### HyperLOPIT Subcellular Fractionation

Subcellular particles were resolved on an iodixanol density gradient as previously described (Christoforou et al. 2016; Mulvey et al. 2017). Briefly, crude subcellular material was enriched by ultracentrifugation over a two-step iodixanol cushion (6% and 25% [w/v] in homogenisation buffer) for 1.5 h at 100,000 × g at 4 °C (SW40Ti rotor, Beckman Optima L-80XP) using maximum acceleration and minimal braking. The supernatant, containing soluble cytoplasmic proteins, was removed, and the visible interphase bands enriched in subcellular particles were collected.

The recovered material was adjusted to 30% (w/v) iodixanol using 50% iodixanol in homogenisation buffer and underlaid beneath a linear gradient prepared from equal volumes of 10% and 20% (w/v) iodixanol solutions that had been allowed to diffuse at 4 °C for 6 h. Gradients were centrifuged for 12 h at 100,000 × g at 4 °C (VTi65.1 rotor, Beckman) with maximum acceleration and minimal braking to separate membranes, organelles and other subcellular particles. Gradients were fractionated into 22 equal volumes by puncturing the tube base and collecting eluate dropwise under gravity. Aliquots were used to determine fraction density (refractive index; Eclipse Handheld Refractometer 45-02, Bellingham + Stanley) and protein concentration (BCA assay, Thermo Fisher Scientific).

### Proteomic Sample Generation

Gradient fractions, together with an aliquot of the cytosolic fraction, were adjusted to equal volumes with water, and the pH raised with 0.1 M triethylammonium bicarbonate (TEAB, Sigma-Aldrich). Triton X-100 was added to 0.1% (v/v) and incubated at room temperature for 10 min to lyse vesicles or intact organelles. Proteins were reduced with 10 mM dithiothreitol for 1 h at 56 °C and alkylated with 25 mM iodoacetamide for 30 min in the dark.

Proteins were precipitated by adding trichloroacetic acid to 10% (w/v) and incubating on ice for 20 min, followed by centrifugation at 18,000 × g for 30 min. Pellets were washed with 10% TCA, centrifuged again and then pellets were resuspended in 500 µl acetone, disrupted by sonication (5 × 30 s on/off cycles, high power) and stored overnight at −20 °C. After centrifugation, acetone was removed, and pellets were air-dried and resuspended in 8 M urea, 100 mM TEAB at ∼1 mg ml−^1^. Protein concentration was determined by BCA assay.

Sequential gradient fractions were pooled into ten groups (20–100 µg protein each). An additional fraction was prepared from the soluble protein pool. Proteins from each pool were digested with Trypsin/Lys-C mix (Promega) at a 1:25 enzyme-to-protein ratio for 4 h at 37 °C, diluted eightfold with 100 mM TEAB and incubated overnight at 37 °C.

After digestion, samples were centrifuged (18,000 × g, 10 min, 4 °C) to remove insoluble material. Supernatants were transferred to LoBind tubes (Eppendorf) and labelled using TMT11plex reagents (Thermo Fisher Scientific) following the manufacturer’s protocol. Briefly, TMT reagents (0.8 mg) were reconstituted in LC-MS grade acetonitrile, mixed with peptide digests and incubated at room temperature for 1 h with shaking (800 rpm). Labelling was quenched with 0.3% (v/v) hydroxylamine for 15 min. All TMT-labelled samples were pooled and dried by vacuum centrifugation. Table S1 summarises the contents of the TMT fractions for all three experiments.

Combined samples were desalted using peptide desalting spin columns (Thermo Fisher Scientific) and eluted in 50% (v/v) acetonitrile, 0.1% (v/v) trifluoroacetic acid prior to final drying by vacuum centrifugation.

### Peptide fractionation

The peptides were fractionated by high-pH reverse-phase chromatography as previously described (Nightingale et al. 2018) using an Ultimate 3000 ultra-high-performance chromatography (UHPLC) system (Thermo Scientific) equipped with a Kinetex® Evo C18 (1.7 μm, 2.1 x 150 mm) column (Phenomenex). The mobile phase used were: LCMS-grade acetonitrile: HPLC-grade water 3:97(v/v) (eluent A); 100% LCMS-grade acetonitrile (eluent B) and 200 mM ammonium formate in HPLC-grade water, pH 10 (eluent C). Eluent C was maintained at 10%, while A and B were altered over the fractionation gradient. The elution gradient was 0-19% B in 10 min; 19-34%B in 14.25 min and 34-50% B in 8.75 min, followed by a 10 min wash of 90 % B. Flow was set at 0.20 ml min-1 for the first 5 min while peptides which were reconstituted in 40 µL of eluent C were loaded onto the column. Thereafter the flow rate was increased to 0.40 ml min-1 for the rest of the protocol. A total of 168 fractions were collected, every 15 seconds between 10 and 52 min of the fractionation run. The fractions were concatenated into 12 pools and reduced to dryness in a vacuum centrifuge.

### Mass Spectrometry

All mass spectrometry analyses were performed on an Orbitrap Fusion™ Lumos™ Tribrid™ instrument coupled to a Dionex Ultimate™ 3000 RSLCnano system (Thermo Fisher Scientific) as previously described (Geladaki et al. 2019), with slight modifications. Briefly, each of the fractionated samples was resuspended in 20 μL of 3% acetonitrile, 0.1% (v/v) aqueous solution of trifluoroacetic acid. Approximately 1 μg of peptides was loaded per injection for LC-MS/MS analysis.

The nano-flow liquid chromatography method for LC-MS/MS was set as follows. Eluent A was 0.1% (v/v) formic acid solution in water. Eluent B was 80% (v/v) aqueous acetonitrile supplemented with formic acid to a final concentration of 0.1% (v/v). The sample loading solvent was 0.1% (v/v) formic acid in water. All solvents and reagents were of HPLC gradient grade or better. Peptides were loaded onto a micro precolumn (100 μm i.d. × 2 cm, particles were C18 PepMap 100, 5 -μm particle size, 100 Å pore size, Thermo Fisher Scientific) using the loading pump for 5 min. After this, the valve was switched from load to inject. Peptides were separated on an EASY-Spray column (PepMap RSLC C18, 50 cm × 75 μm i.d., 2 μm particle size, 100 Å pore size, Thermo Fisher Scientific) using a 7-37% (v/v) gradient of acetonitrile supplemented with 0.1% (v/v) formic acid at 300 nL min^−1^ over 180 min. A wash step (95% Eluent B for 4 min) was included, followed by re-equilibration into Eluent A. The total run time was 210 min.

The MS workflow parameters were set as follows using the Method Editor in XCalibur v4.2.47 (Thermo Fisher Scientific) for the SPS-MS^3^ acquisition method. Detector type: Orbitrap; Resolution: 120,000; Mass range: Normal; Use quadrupole isolation: Yes; Scan range: 380-1,500; RF lens: 30%; AGC target: 4e5; Max inject time: 50 ms; Microscans: 1; Data type: Profile; Polarity: Positive; Monoisotopic peak determination: Peptide; Include charge state(s): 2-7; Exclude after n times: 1; Exclusion duration (s): 70; Mass tolerance (p.p.m.): Low: 10; high: 10; Perform dependent scan on single charge state per precursor only: Yes; Intensity threshold: 5.0e3; Data-dependent mode: Top speed; Number of scan event types: 1; Scan event type 1: No condition; MSn level: 2; Isolation mode: Quadrupole; Isolation window (*m/z*): 0.7; Activation type: CID; CID collision energy (%): 35; Activation Q: 0.25; Detector type: Ion trap; Scan range mode: Auto; *m/z*: Normal; Ion trap scan rate: Turbo; AGC target: 1.0e4; Max inject time (ms): 50; Microscans: 1; Data type: Centroid; Mass range: 400-1200; Exclusion mass width: *m/z*: Low: 18; high: 5; Reagent: TMT; Precursor priority: Most intense; Scan event type 1: No condition; Synchronous precursor selection: Yes; Number of precursors: 10; MS isolation window: 0.7; Activation type: HCD; HCD collision energy (%): 65; Detector type: Orbitrap; Scan range mode: Define *m/z* range; Orbitrap resolution: 60,000; Scan range (*m/z*): 100-500; AGC target: 1.0e5; Max inject time (ms): 120; Microscans: 1; Data type: Profile; AGC, automatic gain control; HCD, higher-energy collisional dissociation; CID, collision-induced dissociation.

An electrospray voltage of 1.5 kV was applied to the eluent via the electrode of the EASY-Spray column. The mass spectrometer was operated in positive ion data-dependent mode for SPS-MS^3^. The total run time was 210 min.

### Quantification and Statistical Analysis

#### Raw LC-MS Data Processing and Quantification

Raw LC–MS data files were processed in Proteome Discoverer v3.1 (Thermo Fisher Scientific) using the Sequest HT server. Searches were performed against the annotated *Plasmodium falciparum* strain 3D7 reference proteome (PlasmoDB release 68, downloaded 25 November 2024) (Aurrecoechea et al. 2009) supplemented with a list of common contaminants (Frankenfield et al. 2022). The SwissProt *Homo sapiens* proteome (UniProt 2025) was also included for quantification but subsequently excluded from downstream analyses.

The Sequest HT search parameters were: precursor mass tolerance 10 ppm, fragment mass tolerance 0.6 Da, trypsin specificity (up to two missed cleavages), fixed modifications (carbamidomethylation of cysteine; TMT 11-plex on peptide N-termini and lysine residues) and variable modifications (methionine oxidation; asparagine and glutamine deamidation; protein N-terminal acetylation and methionine loss). Peptide-spectrum matches (PSMs) were rescored using Inferys (Zolg et al. 2021) (version bundled with PD 3.1), followed by validation with the Percolator-based Peptide Validator node at 1% (strict) and 5% (relaxed) false discovery rate (FDR). Protein identifications were filtered to require at least one unique peptide and validated using the Protein FDR Validator node at 1% (strict) and 5% (relaxed) FDR, applying strict parsimony for protein grouping.

Reporter ion intensities (TMT 11-plex) were quantified using the Reporter Ions Quantifier node, with quantification restricted to unique peptides, a co-isolation threshold of 75%, an SPS mass match threshold of 65%, and a minimum average reporter signal-to-noise of 10. Iterative isotopic impurity correction was applied, and no further normalisation or scaling was performed. Quantification results were exported as tab-delimited text files and processed in R (v ≥ 4.5) using MSnbase (v ≥ 2.34.0) and pRoloc (v1.48.0) from Bioconductor, together with tidyverse (v 2.0.0) packages.

For quantification, the PSM-level output was filtered to retain only unique, unambiguous matches assigned to a single protein group (excluding contaminants), with rank 1 peptides, ≤50% precursor isolation interference, ≥50% matched SPS precursor ions and an average reporter ion signal-to-noise ratio ≥ 5. PSMs with missing reporter ion values were removed if alternative PSMs with complete quantitation existed for the same protein group, or if more than four values in the middle eight TMT channels (TMT127C–TMT130N) were missing. Remaining missing values were imputed using the hot-deck ‘knn’ method implemented in MSnbase::impute. Prior to imputation, reporter ion intensities were normalised to their sum across channels to reduce intensity variance. Row-sum-normalised abundances for each protein group were summarised to protein-level values using median TMT reporter intensities. Proteins with complete abundance profiles across both 10plex datasets were retained, and contaminants and non-*Plasmodium* proteins were excluded from downstream analysis.

### Data Analysis and Protein Localisation Assignment

The PSM-level output from Proteome Discoverer was imported into R and analysed using the MSnBase and pRoloc packages from Bioconductor (Gatto et al. 2014; Gatto and Lilley 2012; Gentleman et al. 2004). Aggregation of PSMs into peptides and subsequently proteins was performed as previously described (Guerin et al. 2023). The protein data from three experiments was concatenated to form a single dataset with 2443 proteins containing 30 quantified samples. t-distributed Stochastic Neighbour Embedding (t-SNE) (Van der Maaten and Hinton 2008) was used for reducing the dimensions of the data to produce a visible, two dimensional plot.

To perform t-SNE, the data was PCA-transformed and the top principal components that accounted for 99% of the variance were kept. These principal components were filtered and subjected to a statistical whitening transformation such that each principal component had variance 1. Embedding into two dimensions was performed, using a perplexity value of 50 for a maximum of 10,000 iterations. This output produced x and y co-ordinates for each protein, which were recorded and used to produce a two-dimensional map.

For supervised machine learning classification of protein locations, a set of 542 manually curated marker proteins defining 24 subcellular classes was compiled using previously published localisation data, or information available from public sources such as gene annotation in PlasmoDB.org or the ApiLoc database (Woodcroft et al. 2011).

Subcellular protein localization was inferred using supervised machine learning with Support Vector Machines (SVMs) deployed through the pRoloc package in R (Breckels et al. 2016) using a radial basis function kernal. Our marker proteins were used to train the classifier. To optimize the performance of the SVM model, a grid search was conducted across multiple combinations of cost and sigma parameters, repeated over 100 iterations. The optimal parameter set was selected based on the highest mean F1 score across these cross-validation rounds. Following optimization, the final model was trained on the full marker set and SVM localization scores were generated for all proteins in the dataset. These scores were then converted to class labels by thresholding on the median SVM score per protein. Furthermore, to improve classification specificity, particularly for organelles where overfitting was observed, thresholds for several subcellular classes were manually refined based on empirical score distributions and existing biological knowledge. Proteins above the threshold were assigned to a subcellular compartment and those below the threshold were left “unlabelled”.

### Validation of Protein Localisation

To validate predicted protein localisations, we applied a Selection-Linked Integration (SLI) approach to target gene loci. In this method, ∼800 bp of the 3′ coding region of the gene of interest was PCR-amplified and cloned into a targeting plasmid designed to fuse a C-terminal tag to the gene of interest following selection for integration (Jonsdottir et al. 2021) using Gibson assembly. Oligonucleotide primers used in this study are listed in Table S3.

For transfection, synchronous *P. falciparum* cultures at 5% parasitaemia were electroporated with 60 µg purified plasmid DNA using a Bio-Rad Gene Pulser (310 V, 950 µF). Following transfection, parasites were maintained under 5 nM WR99210 selection (Jacobus Pharmaceuticals) until resistant populations emerged. For SLI, WR99210-resistant parasites were subsequently treated with G418 (Sigma-Aldrich) to select for integrants as described previously (Birnbaum et al. 2017).

Successful genome editing was confirmed by PCR amplification of genomic DNA. Tagged proteins were detected by Western blot using an anti-HA antibody (Roche, ROAHAHA).

For immunofluorescence microscopy, parasites were either fixed directly or first stained live with MitoTracker™ prior to fixation. For mitochondrial labelling, schizont-stage parasites were incubated with 100 nM MitoTracker™ Red CMXRos (Invitrogen) in Albumax-free culture medium for 30 min at 37 °C, then washed once in pre-warmed RPMI medium before fixation. Cells were fixed either in 4% paraformaldehyde (PFA) and 0.0075% glutaraldehyde (GA) in phosphate-buffered saline (PBS) for 20 min at room temperature, followed by three PBS washes (Tonkin et al. 2004). Cells were permeabilised with 0.1% Triton X-100 in PBS for 10 min and blocked with 3% bovine serum albumin (BSA) in PBS for 1 h at room temperature. Primary antibodies (Table S3) diluted in 3% BSA/PBS were incubated with cells for 1 h, followed by three PBS washes. Alexa Fluor-conjugated secondary antibodies were applied for 1 h in the dark, followed by additional PBS washes. Cells were mounted in Fluoromount-G™ Mounting Medium, with DAPI (Invitrogen, 00-4959-52) before imaging.

Images of ER, PM and host PM proteins were acquired using a Nikon Eclipse Ti wide-field microscope with a Nikon objective lens (Plan APO, 100×/1.45 oil) and a Hamamatsu C11440, ORCA Flash 4.0 camera. Images of IMC, mitochondrion and rhoptry proteins were acquired using a Leica SP8 confocal microscope (operated with its Leica LAS X interface) with a 63x lens (1.4 oil) and a combination of two PMT fluorescence detectors and one hybrid GaAsP detector (HyD). All fluorescence images were processed using Fiji software (https://imagej.net/software/fiji/). The antibodies used, source, and the relevant concentrations are described in Table S3.

### MalariaGen data analysis

VCF files containing genomic variation data from 20,864 *P. falciparum* isolates from 33 countries were downloaded from the Pf dataset v7.0 hosted in MalariaGEN (MalariaGen et al. 2023). VCF processing was carried out using bcftools v1.11 (Danecek et al. 2021) as follows: Multiallelic sites were normalised, keeping only single nucleotide variations with allele frequencies higher than 0.01 and lower than 0.99. Potential duplicates were removed, and the annotation field was removed. Variant effects were predicted with SnpEff v.5.21f (Cingolani 2022) using a custom database built from the annotations deposited in PlasmoDB release 68 (Harb, McDowell, and Roos 2024). We calculated the variant fraction per gene as the number of observed variant sites divided by the length of the coding region; the adjusted proportion of non-synonymous over synonymous variants (adj. pN/pS) was calculated by dividing the number of non-synonymous variants by the sum of synonymous and non-synonymous variants (supplementary table S2 and S4).

### GO enrichment analysis

Gene Ontology (GO) terms were downloaded from PlasmoDB release 68, GO enrichment analysis was performed using the enricher function from clusterProfiler v4.16.0 (Xu et al. 2024) with the GO terms of the proteins present in each location against the GO terms in the rest of the proteins detected in the LOPIT experiments. p-values were adjusted using Benjamini-Hochberg correction. GO terms with an adjusted p-value smaller than 0.01 were considered significant.

### Orthology analysis

The coding sequences of 21 organisms (table S5) were used to detect clusters of orthologous genes. The amino acid sequences were preprocessed with OrthoPrep v.0.0.1 (https://github.com/vflorelo/orthoprep), reciprocal best hits were detected using diamond with an e-value of 1e-6, the processed sequences were further analysed with OrthoFinder v2.5.5 (Emms and Kelly 2019), clusters of orthologous genes were detected with mcl v14.137 using an inflation coefficient of 1.1.

### Gene gains at different divergence points

Gene counts per species for each orthogroup (OG) were obtained from the Orthofinder output; they were transformed into a presence/absence matrix. Orthogroup gains across the evolutionary history of *P. falciparum* were calculated with the ete3 toolkit (Huerta-Cepas, Serra, and Bork 2016) using a consensus species tree as a reference. The species tree was built based on recent analyses (Munoz-Gomez et al. 2019; Galen et al. 2018; Janouskovec et al. 2019). Orthogroup gains consisted of OGs that were present in the outgroup of a divergence point and at least one more descendant.

#### Evolutionary enrichment of subcellular compartments

Using the consensus species tree, compartment-specific enrichments at nine points of divergence in the evolutionary history of *P. falciparum* using hypergeometric tests taking the total number of localised proteins as the background distribution, and calculating the significance of detecting the number of proteins per location using the number of gained proteins at each divergence point with a confidence interval of 95%. Benjamini - Hochberg p-value correction was performed for false discovery rate. Intersections of subcellular locations and divergence points were considered significantly enriched if their q-value was lower than 0.05 and the number of proteins was higher than that expected from chance.

### Distribution of evolutionary selection pressures

Using orthologous genes identified by OrthoFinder (see above), codon-based multiple sequence alignments were constructed using mafft v7.475 (Nakamura et al. 2018) and pal2nal (Suyama, Torrents, and Bork 2006). Phylogenetic trees for each group of orthologous genes were built with iqtree2 (Minh et al. 2020) from the amino-acid sequence alignments. Each phylogenetic tree was used as a reference for the maximum likelihood estimation of dN/dS ratios from the corresponding codon-based multiple nucleotide sequence alignments using PAML v4.10.6 (Yang 2007). Average dN/dS ratios were calculated from the output of PAML-codeml.

### Disorder prediction

Disordered protein regions were assigned using four different sources. Disordered region predictions were made using iupred3 (Erdos, Pajkos, and Dosztanyi 2021) and MobiDB-lite v3.9.0 (Necci et al. 2017), in both cases, we only kept predictions longer than 30 amino acids. AlphaFold predicted 3D structures of each *P. falciparum* 3D7 protein were taken from Uniprot (proteome accession number: UP000001450). The secondary structure elements were parsed from the PDB files using DSSP v.2.3.0 (Kabsch and Sander 1983), low confidence structural elements were parsed manually from the PDB files. Any string with more than 30 amino acid residues with either low confidence prediction, or, lack of secondary structure features, was considered a disordered region. Once we had the predictions of disordered regions from four different sources (iupred3, MobiDB-lite, AlphaFold+DSSP, AlphaFold LowConf), we built consensus predictions corresponding to regions that were predicted as disordered by at least two sources, the same threshold of 30 amino acids was applied for keeping consensus disordered regions. We calculated the disordered fraction by dividing the cumulative length of the predictions (disordered length) by the total length of each protein.

### Data integration and statistical analysis

We compared the distributions of seven metrics (Figures 5-7, supplementary table S2), across the different subcellular compartments to test if there were significant patterns between the evaluated metric and each subcellular compartment. At each compartment we conducted a two-sample Kolmogorov-Smirnov test (KS) to compare the distribution of each metric of proteins in a location against proteins in the remaining compartments. Compartments for which there were less than 20 data-points were not subjected to statistical testing. Metric distributions were considered significantly different if their KS p-value was less than 0.05.

## SUPPLEMENTARY TABLES AND FIGURES

Table S1: Summary of gradient fractions and pooling strategies for mass spectrometry

Table S2: Summary of classifications and findings from this study

Table S3: List of oligonucleotide primers, antibodies and other reagents used in this study.

Table S4: Full calculations of MalariaGen analysis

Table S5: Genomes used for orthology analyses.

**Figure S1.**
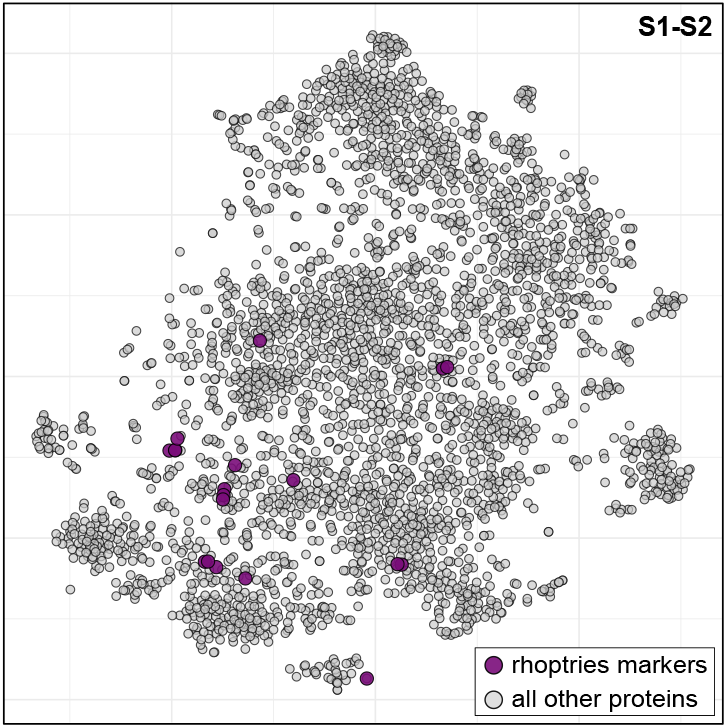
Rhoptry markers do not cluster in the S1-S2 combined analysis. t-SNE projection of 3000 proteins from the S1-S2 concatenated data with rhoptry marker proteins shown in purple.

**Figure S2.**
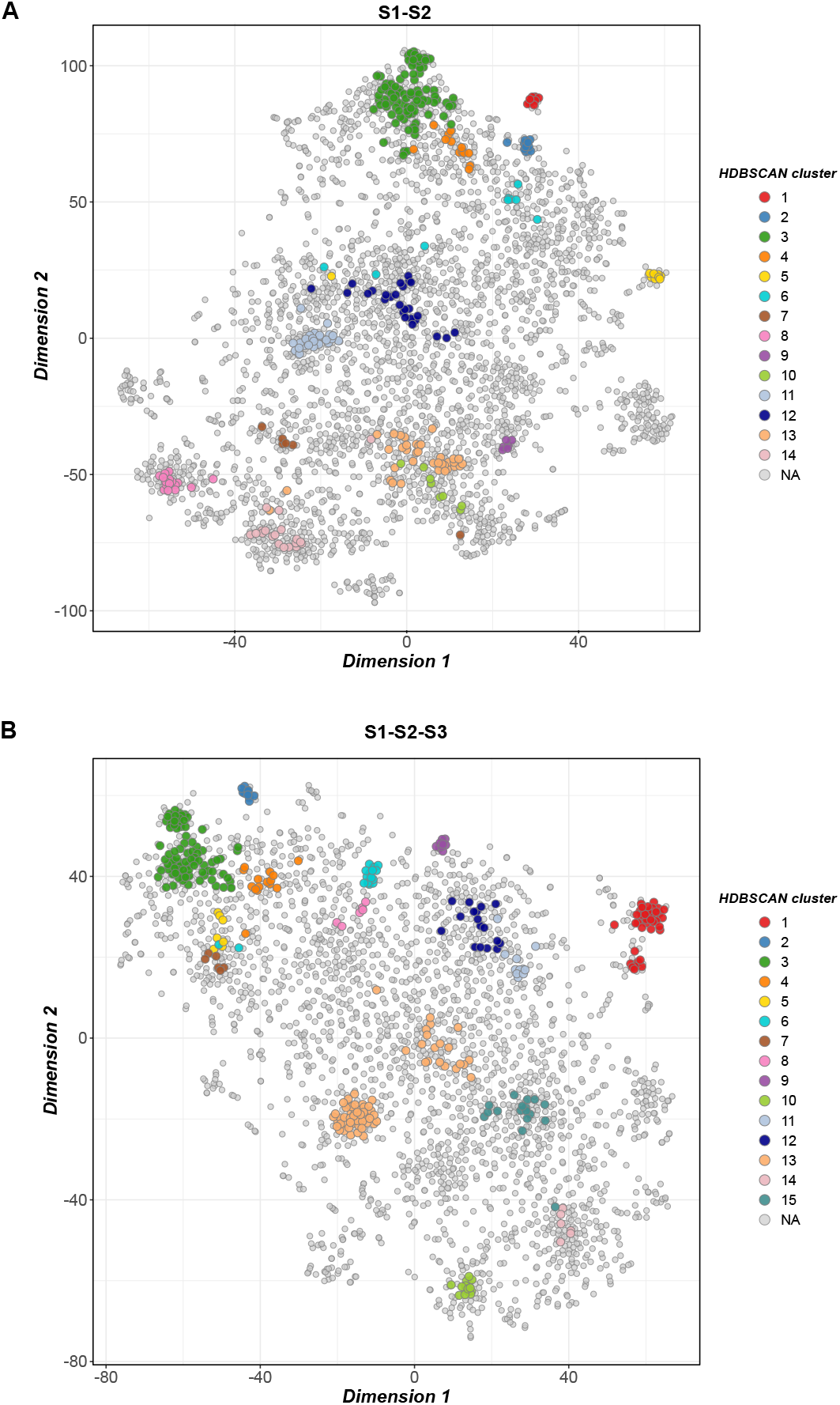
Unsupervised clustering using HDBSCAN. Protein clusters discovered by analysis of raw abundance-distribution profiles of the S1-S2 dataset (**A**) and S1-S2-S3 dataset (**B**) with HDBSCAN overlaid on the t-SNE projection. Distinct clusters are numbered and indicated by colour. Core distance was set to the 8th nearest neighbour (min_samples = 8) with a minimum cluster size of 8 (min_cluster_size = 8). Clusters were defined using the ‘leaf’ method; all other parameters were default.

**Figure S3.**
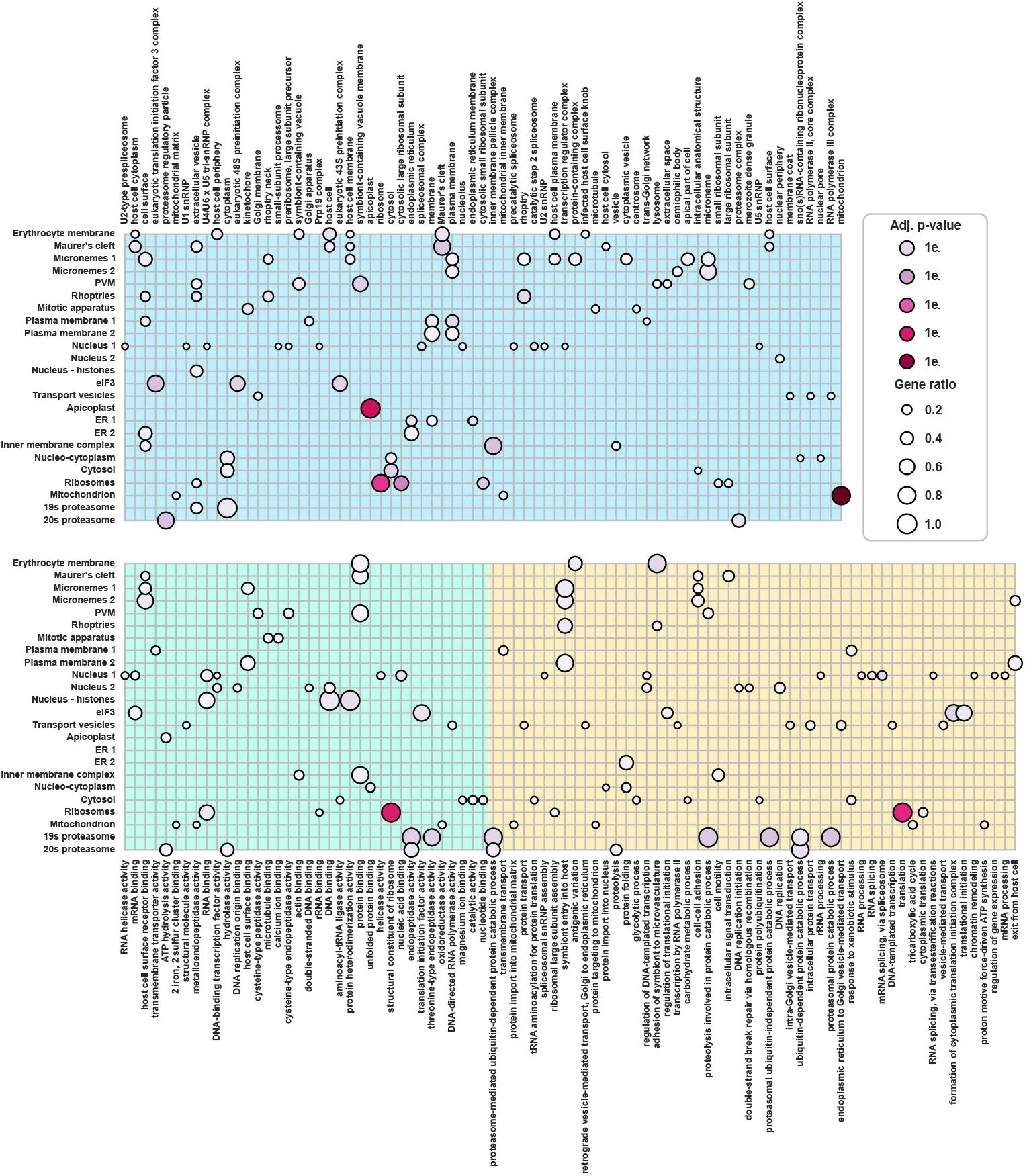
GO enrichment analysis of organelle proteomes. Bubble plot showing the results of GO enrichment analysis. GO terms (x-axis) from the proteins in each compartment (y-axis) were analysed with clusterProfiler. Only significant terms (adjusted p-value smaller ≤ 0.01) are shown, size of the dots represents the proportion of proteins showing a specific term against the total number of proteins in each compartment. GO categories are shown as follows: Cellular Component in blue, Molecular Function in green and Biological Process in yellow

**Figure S4.**
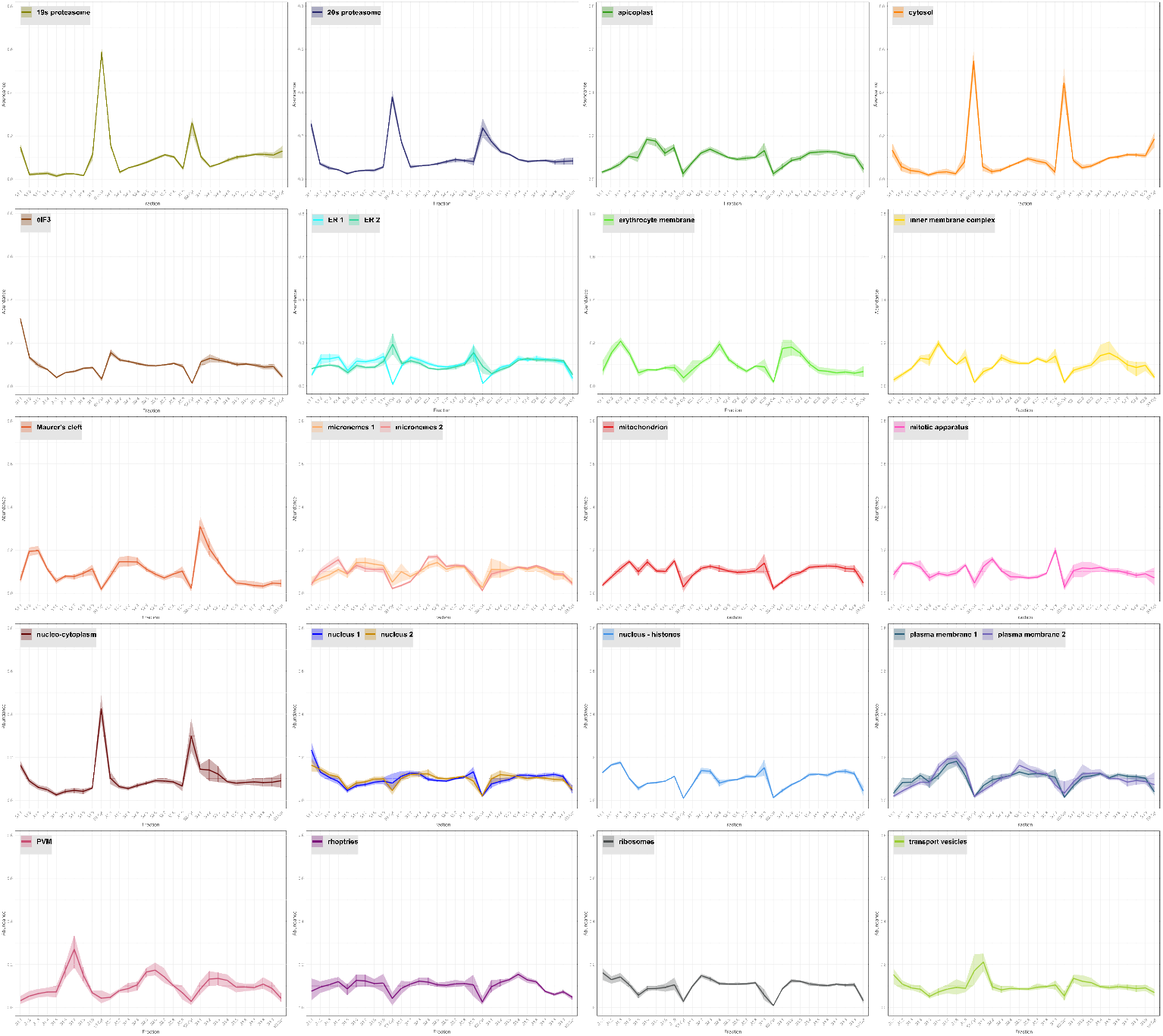
Abundance distribution profiles of organelle marker proteins. Normalised abundance profiles are shown for marker proteins across three independent hyperLOPIT experiments (S1-S2-S3), each analysed by TMT 10-plex labelling. In each experiment, nine fractions from an iodixanol density gradient (S1.1–S1.9, S2.1–S2.9 and S3.1– S3.9) and a cytosolic fraction (S1.Cyt, S2.Cyt and S3.Cyt) were each labelled with an individual TMT channel. Profiles from the three experiments were concatenated to generate a combined dataset. Organelles with subclusters are shown in individual plots, with the subclusters highlighted in different colours.

**Figure S5.**
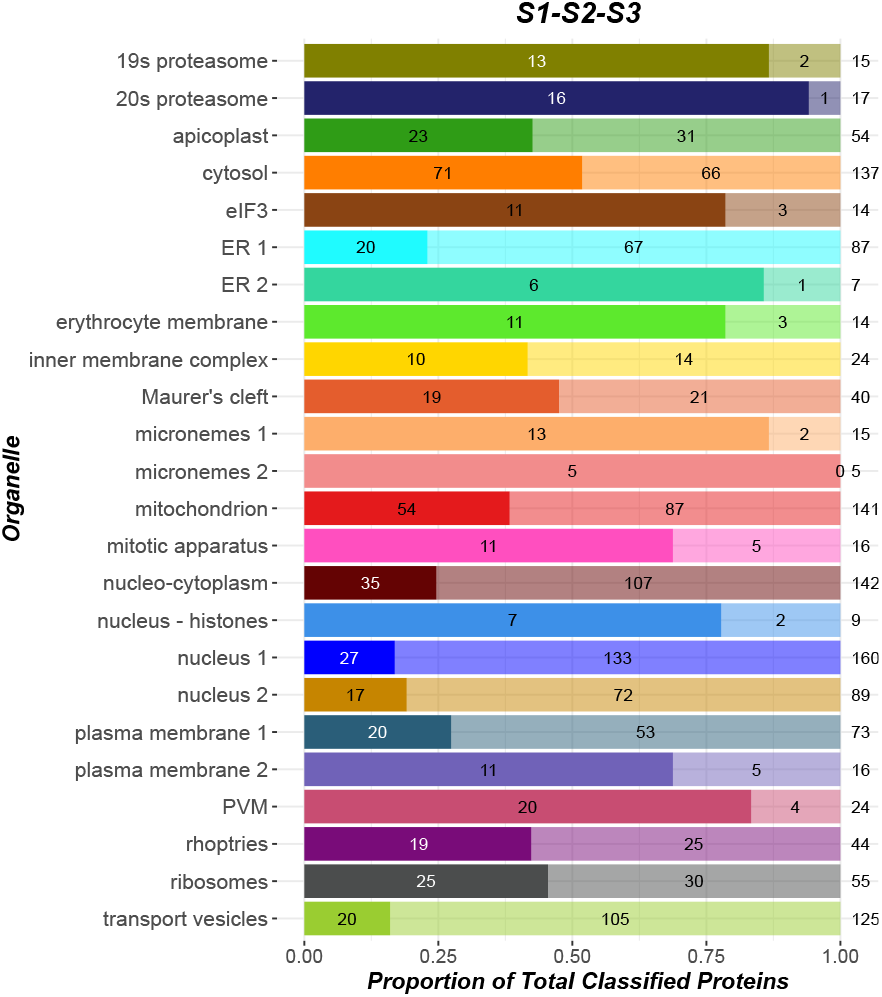
Results of supervised machine learning on S1-S2-S3. Numbers of proteins assigned to each compartment by SVM classification from the S1-S2-S3 analysis shown as bars with markers numbers to the left and predicted proteins to the right of the bars (totals at far right.

**Figure S6.**
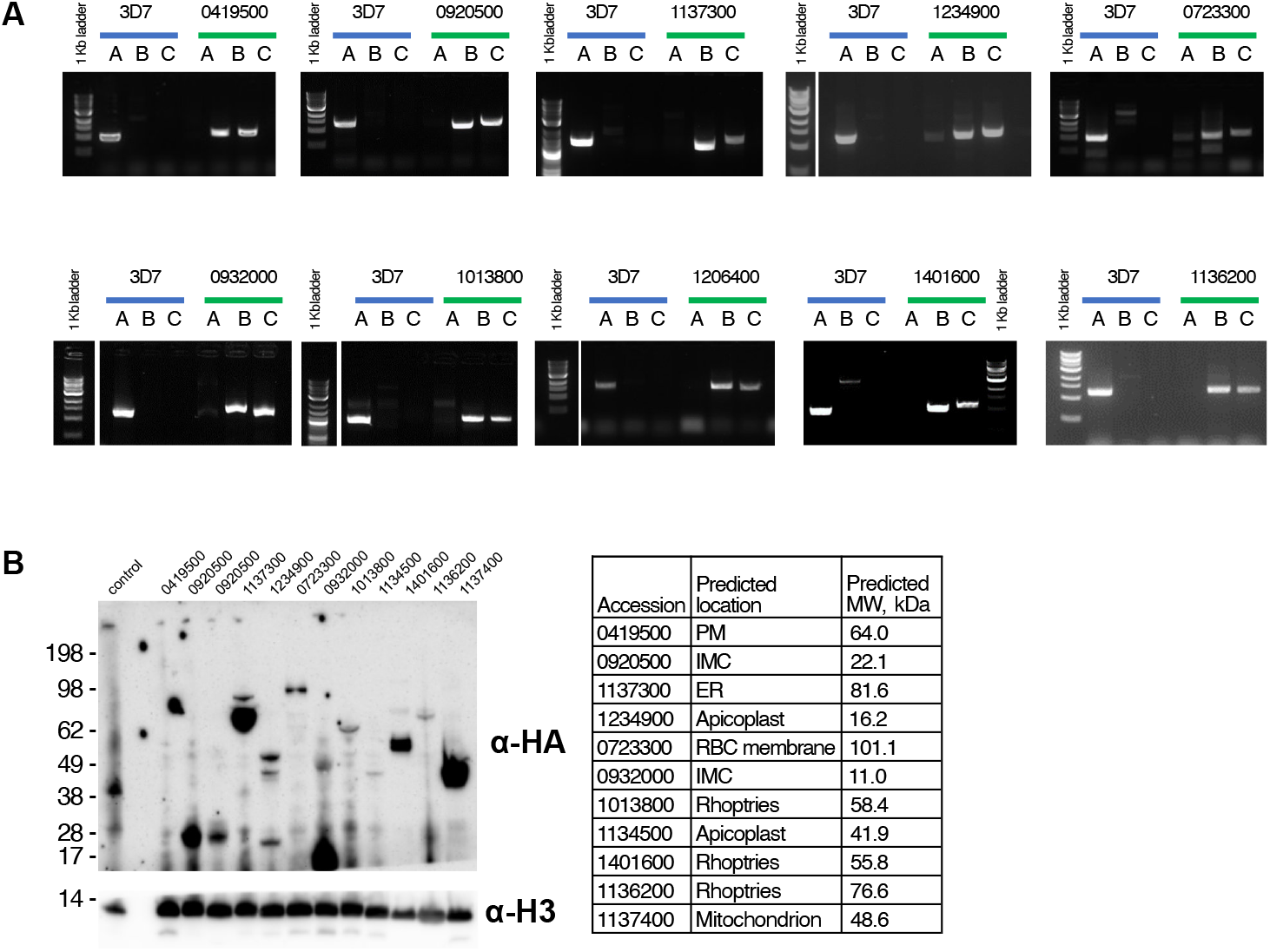
Validation of C-terminal 3×HA tagging and genomic integration of candidate proteins in *Plasmodium falciparum*. **A**) Diagnostic PCRs confirming correct integration of the tagging cassette at the endogenous locus for each target gene. Lane A tests amplification of the wild-type (WT) locus, lane B tests 5′ integration, and lane C tests 3′ integration. Successful integration is indicated by amplification of both 5′ and 3′ integration products and loss of the WT band. **B**) Western blot of schizont-stage parasite lysates from individual, uncloned transfectant lines probed with anti-HA antibody, confirming expression of C-terminally 3×HA-tagged proteins. IDs correspond to PlasmoDB identifiers (3D7 strain). Anti-Histone H3 was used as a loading control. The predicted molecular weights of each tagged protein are listed in the adjacent table. Two proteins (1234900 and 1134500) produced a band of the expected size by western blot but were below the detection limit in fixed-cell IFA imaging, possibly due to low expression levels or epitope masking so are not reported on further here.

## Notes

### Competing Interest Statement

The authors have declared no competing interest.

